# Unlocking Trypanosome Biology: A Comprehensive Protein-Tagging Toolkit for Localization and Functional Analysis

**DOI:** 10.1101/2023.04.21.537815

**Authors:** Athina Paterou, Jiří Týč, Jack Sunter, Sue Vaughan, Keith Gull, Samuel Dean

## Abstract

African trypanosomes are medically important parasites that cause Sleeping sickness in humans and nagana in animals. In addition to their pathogenic role, they have emerged as valuable model organisms for studying fundamental biological processes. Protein tagging is a powerful tool for investigating protein localization and function. In a previous study, we developed two plasmids for rapid and reproducible protein tagging in trypanosomes, which enabled the localisation of all proteins in the trypanosome cell. However, the limited selection of fluorescent protein tags and selectable markers restricted the flexibility of this approach. Here, we present an expanded set of >100 vectors that utilizes universal primer annealing sequences, enabling protein tagging with a range of fluorescent and biochemical tags using five different selection markers. We evaluated the suitability of various fluorescent proteins for live cell imaging and determined their brightness and stability under different fixation conditions. Finally, we determined the optimal fluorescent protein for a set of specific experimental conditions demonstrating the utility of this toolkit.

## Introduction

Trypanosoma brucei is a protozoan parasite that causes devastating diseases in humans and animals, including Sleeping sickness in tsetse endemic Africa. In addition to their pathogenic role, trypanosomes have become valuable model organisms for studying fundamental biological processes due to their extreme and tractable biology, including polycistronic transcription, RNA editing, and antigenic variation. Protein tagging is a powerful tool for studying protein localization, interactions, and function [1,2]. Previously, two plasmids were developed for rapid and reproducible protein tagging in trypanosomes [3]. These plasmids utilised a PCR-based approach, whereby a drug resistance cassette and a protein tag were amplified from the plasmid vector, with the amplicon targeted for integration into the gene of interest by the long 5’ overhangs on each primer. Although widely used, a more comprehensive toolkit is needed to enable efficient and versatile protein tagging with a variety of fluorescent, epitope and biochemical tags in different trypanosome strains.

To address this, we developed a set of more than one hundred vectors with universal primer annealing sequences to enable efficient and versatile protein tagging with a variety of fluorescent and biochemical tags in different trypanosome strains. We evaluated the suitability of different fluorescent proteins, including mNeonGreen [4], mGreenLantern [5], and mScarlet-I [6], for live cell imaging and determined their brightness and stability under different fixation conditions. We demonstrated the utility of this toolkit in several different use cases, and our findings provide valuable insights into the suitability of different fluorescent proteins for live cell imaging and time-lapse microscopy. This toolkit will be valuable for investigating protein localization and function in trypanosomes, and will facilitate the study of fundamental biological processes in this important model organism.

## Results

### Design principles and diversity of pPOTv6 and v7 series

Here, we describe a new series of PCR-only Tagging (pPOT) tagging vectors with improved design features, including universal primer annealing sequences to enhance flexibility and reduce primer synthesis costs, and the same drug resistance cassette for both amino and carboxyl terminal tagging (Figure 1). Additionally, these vectors have modules flanked by unique restriction enzyme sites to facilitate their exchange, and offer five different drug resistance genes for tagging in different trypanosome host strains and for generating complex cell lines. The pPOTv6 vectors are designed such that the primary tag is flanked by three copies of an epitope tag (Ty1 [7] or Myc [2]) to facilitate western blot or immunofluorescence analyses, or the 6-His tag to facilitate affinity purification, while the pPOTv7 series vectors have no flanking epitopes. This new vector series encompasses over 20 different fluorescent proteins, three different split fluorescent proteins, three different tags for ‘click’ chemistry, two different proximity labelling tags, and several “fusion tags” allowing microscopy to be combined with another assay. In addition, we designed a set of vectors encoding ten tandem copies of an epitope tag (including FLAG, HA, Myc, Ty, and V5 [2]) to support the emerging technique of expansion microscopy [8–10]. A summary of the complete vector series is presented in Table 1, a comprehensive vector list is in Supplemental Table 1, and all vectors and their GenBank files are available via Addgene (https://www.addgene.org).

**Table 1.**
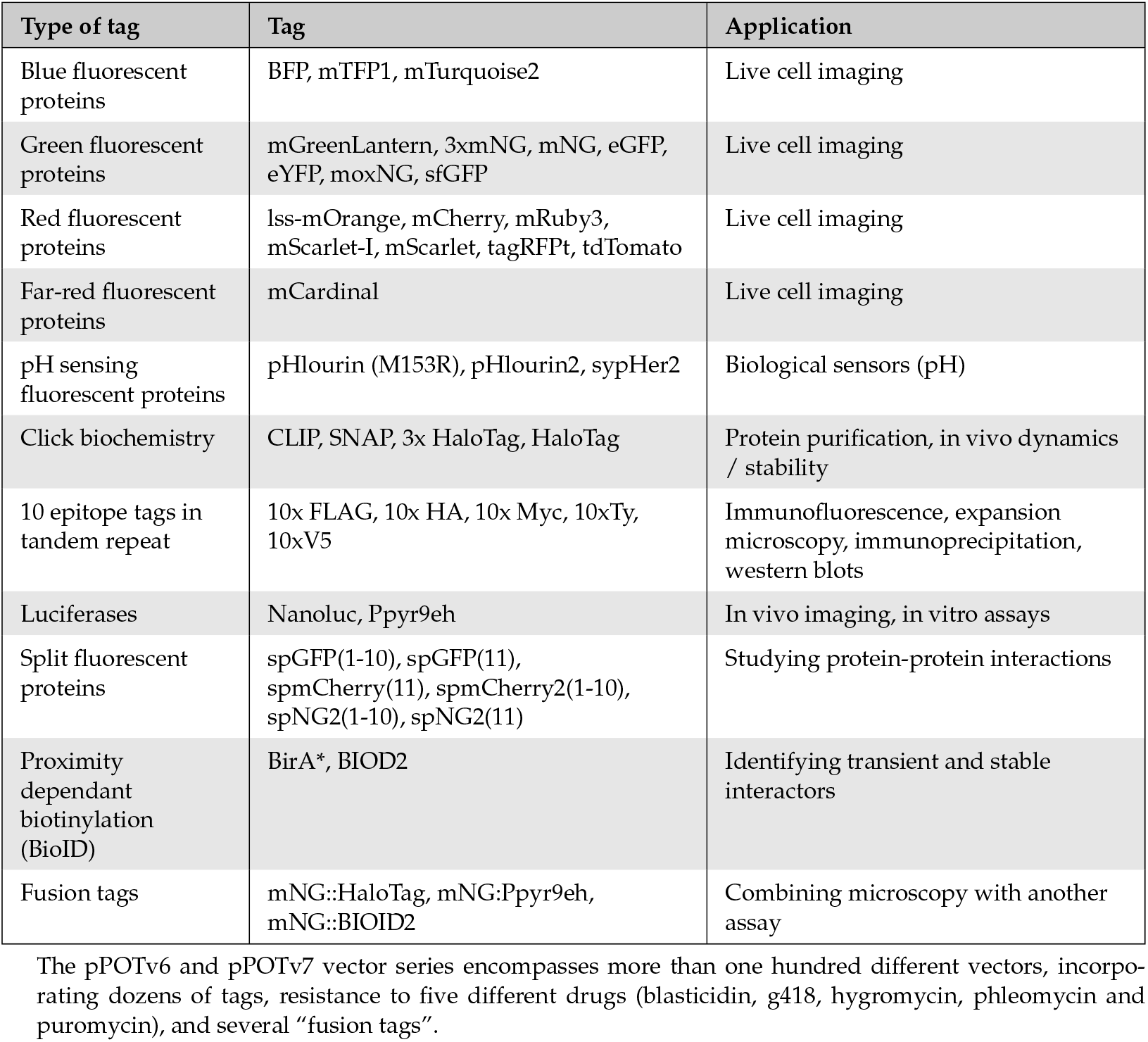
A summary of pPOTv6 and pPOTv7 vectors.

| Type of tag | Tag | Application |
| --- | --- | --- |
| Blue fluorescent proteins | BFP, mTFP1, mTurquoise2 | Live cell imaging |
| Green fluorescent proteins | mGreenLantern, 3xmNG, mNG, eGFP, eYFP, moxNG, sfGFP | Live cell imaging |
| Red fluorescent proteins | lss-mOrange, mCherry, mRuby3, mScarlet-I, mScarlet, tagRFPT, tdTomato | Live cell imaging |
| Far-red fluorescent proteins | mCardinal | Live cell imaging |
| pH sensing fluorescent proteins | pHlourin (M153R), pHlourin2, sypHer2 | Biological sensors (pH) |
| Click chemistry | CLIP, SNAP, 3x HaloTag, HaloTag | Protein purification, in vivo dynamics / stability |
| 10 epitope tags in tandem repeat | 10x FLAG, 10x HA, 10x Myc, 10xTy, 10xV5 | Immunofluorescence, expansion microscopy, immunoprecipitation, western blots |
| Luciferases | Nanoluc, Ppyr9eh | In vivo imaging, in vitro assays |
| Split fluorescent proteins | spGFP(1-10), spGFP(11), spmCherry(11), spmCherry2(1-10), spNG2(1-10), spNG2(11) | Studying protein-protein interactions |
| Proximity dependant biotinylation (BioID) | BirA*, BIOD2 | Identifying transient and stable interactors |
| Fusion tags | mNG::HaloTag, mNG:Ppyr9eh, mNG::BIOID2 | Combining microscopy with another assay |

The pPOTv6 and pPOTv7 vector series encompasses more than one hundred different vectors, incorporating dozens of tags, resistance to five different drugs (blastidicin, g418, hygromycin, phleomycin and puromycin), and several “fusion tags”.

**Figure 1:**
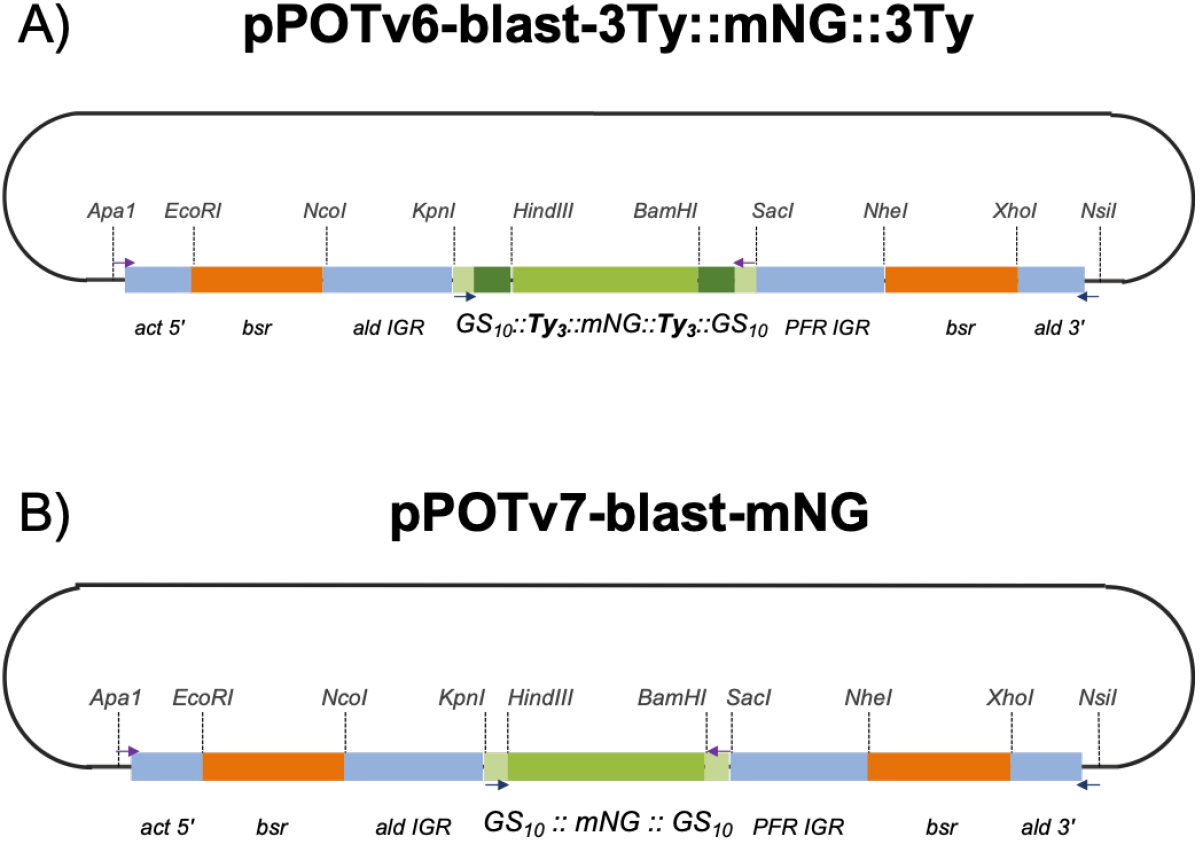
Schematics of pPOTv6 and pPOTv7 design features using A) pPOTv6-blast-3Ty::mNG::3Ty and B) pPOTv7-blast-mNG as examples. Modules such as primary and epitope tags, intergenic sequences and drug resistance genes are flanked by unique restriction enzyme to facilitate their exchange. Primer annealing site used in the generation of tagging amplicons are within sequence encoding GS linkers and common to v6 and v7 series plasmids. Note that tagging using pPOTv6 series vectors results in the tagged protein being fused to the primary tag flanked by three tandem copies of an epitope (six total). Act = actin, ald = aldolase, bsr = blasticidin S-resistance, IGR = intergenic region, PFR = paraflagellar rod, 5’ = 5’ IGR, 3’ = 3’ IGR.

### Brightness of fluorescent protein tags

Most fluorescent proteins have available information on their quantum yield, in vitro brightness and photostability [11]. However, such information may not reflect their behaviour when used as a protein tag in a cell due to the tagged protein’s properties and local environment [12]. To assess their in vivo utility as protein tags, the flagellar Transition Zone Protein 157kD (TZP157) [13] was tagged on the N terminus using 17 different fluorescent proteins. TZP157 was chosen because it forms a small, single focus inside each trypanosome cell that can be easily quantified using custom scripts (Figure 2a, Supplemental Figure 1). The signal intensity in a 1-second exposure was quantified from >100 cells (Figure 3b), and it was found that under these conditions, mNeonGreen was the brightest single copy fluorescent protein tag (Figure 3c). A triple mNeonGreen tag, consisting of 3 copies of mNeonGreen in tandem repeat, was 50% brighter than a single mNeonGreen. Interestingly, mGreenLantern, which was recently published as a viable alternative to mNeonGreen [5], was the least bright of the green fluorescent proteins tested and highlights that fluorescent proteins behave differently in different contexts. tdTomato [14] was the brightest red fluorescent protein, while mScarlet-I [6] was the brightest monomeric red fluorescent protein. mCherry [14], a widely used red fluorescent protein, did not perform well in this assay and showed significant cellular background (Supplemental Figure 1). The only far-red fluorescent protein tested, mCardinal [15], was faint but detectable, suggesting it as an option for making 3-color trypanosomes or deep-tissue imaging of trypanosomes in vivo. Blue fluorescent protein tags did not give a detectable signal at the base of the flagellum (Supplemental Figure 1). To determine whether the high cellular background associated with ultraviolet irradiation prevented their detection, cellular soluble material was extracted using non-ionic detergent to generate cytoskeletons. However, blue-tagged proteins were still not detected, suggesting that they are either too dim or unstable to be useful for most protein tagging in trypanosomes (Supplemental Figure 1).

**Figure 2:**
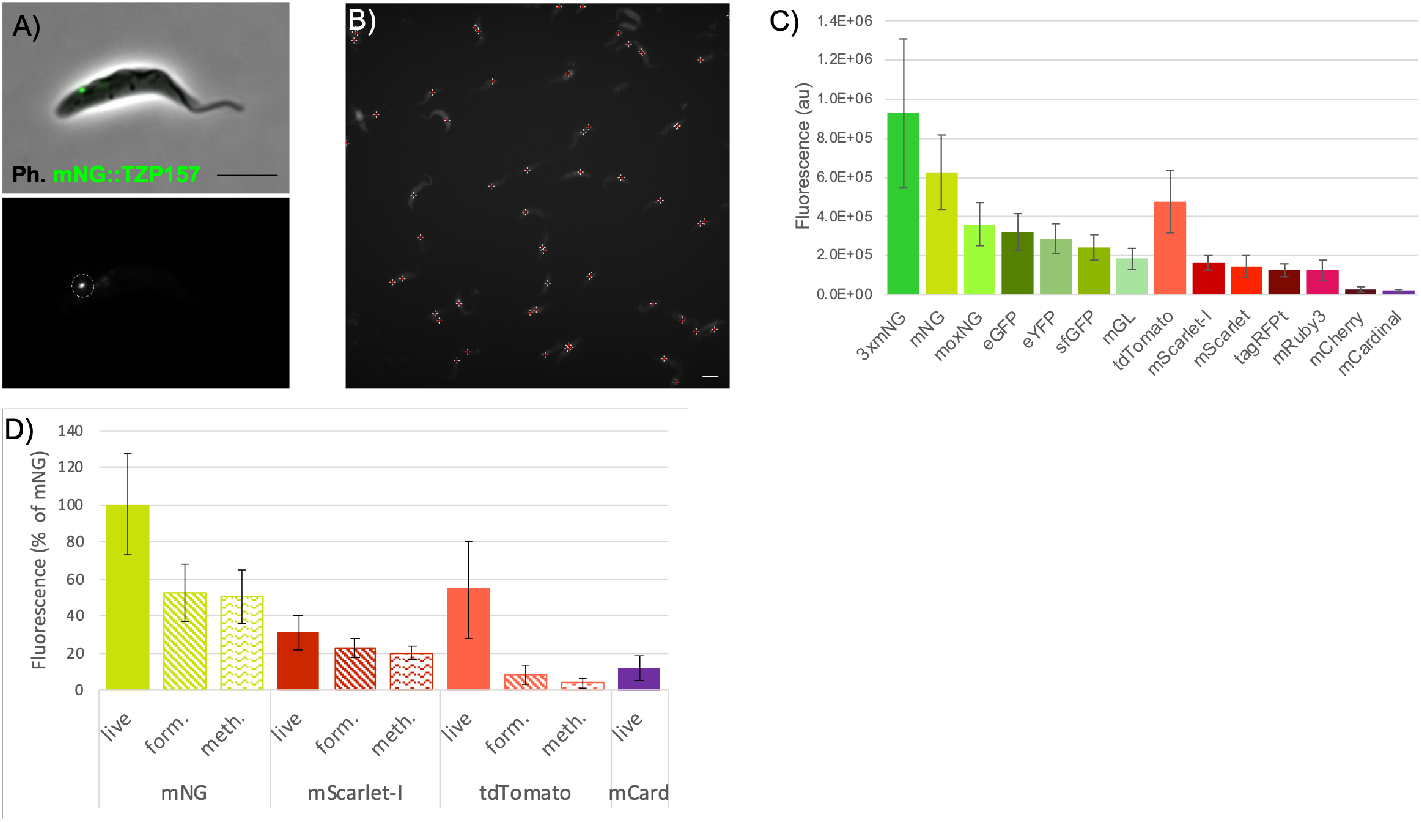
Assessing the brightness of fluorescent proteins tags and their performance in different fixatives. A) An example cell expressing mNeonGreen-tagged TZP157 at the base of the flagellum. B) An example field of cells expressing mNeonGreen-tagged cells with each bright green spot marked for sub-sequent analysis of its brightness. C) The brightness of each fluorescent protein was assessed by quantifying the fluorescent signal emanating from the tagged protein. 3xmNG indicates 3 tandem repeat copies of mNeonGreen and was the brightest tag assessed in this work. D) The effect of formaldehyde and methanol fixation upon the brightness of mNeonGreen, mScarlet-I and tdTomato was assessed by acquiring a 2-second exposure of fixed versus non-fixed cells. Error bars = standard deviation. Scale bar = 5 *μ*m.

**Figure 3:**
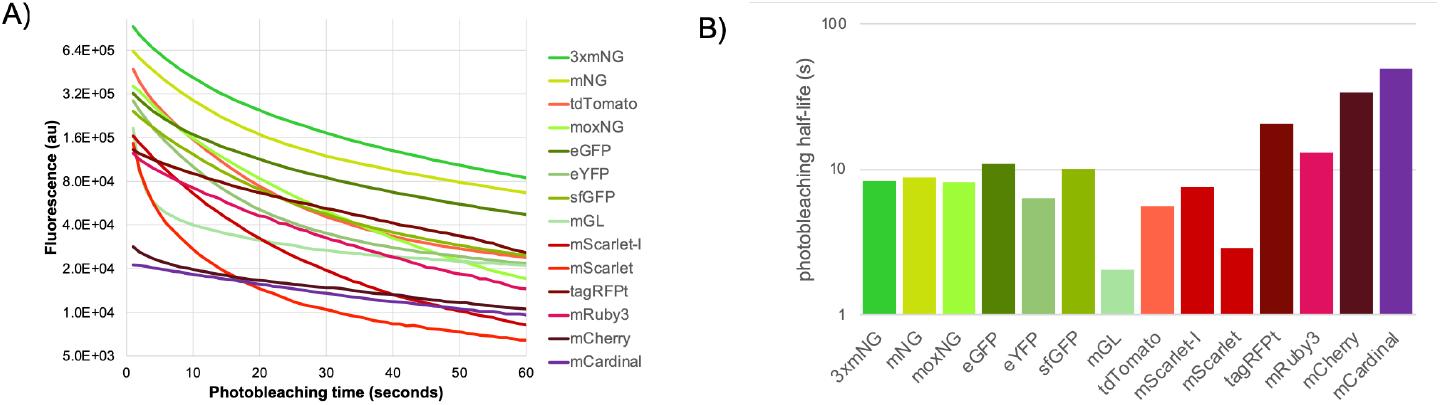
Assessing the photostability of fluorescent protein tags. A) Cells were irradiated over a 30 × 1 second time lapse fluorescence experiment and spots corresponding to tagged TZP157 were quantified to assess the tag’s photostability. B) The photostability half-life was calculated by linear interpolation of the two time points that were closest to half the brightness value after one second.

Based on our experience, the best microscopy data with the fewest artifacts is obtained through live cell imaging. However, in certain situations, such as performing immunofluorescence analyses, it may be necessary to fix cells prior to imaging. To evaluate the performance of fluorescent protein tags under different fixation conditions, a range of the most promising fluorescent proteins fused to TZP157 were imaged for 2 seconds after fixation with formaldehyde or methanol and compared with live, unfixed cells (Supplemental Figure 2). Fixation led to a reduction in the brightness of mNeonGreen fluorescence by approximately 50%, but nonetheless, it was comparable with unfixed tdTomato (Figure 2d). Surprisingly, formaldehyde fixation resulted in a 5-fold reduction in the brightness of tdTomato, and methanol fixation led to a 10-fold reduction. This dramatic decrease in brightness could be attributed to fixatives disrupting the association between individual subunits of the tdTomato dimer. Fixation reduced the brightness of mScarlet-I by only 30-40%, making it a suitable option for imaging fixed red fluorescent proteins. However, the brightness of mCardinal was so greatly reduced by fixation that it was undetectable from cellular autofluorescence (Supplemental Figure 2). Extraction of cytoso-lic material using detergent revealed a spot at the flagellar transition zone after methanol fixation (Supplemental Figure 2), suggesting that in certain cases mCardinal could be a useful tag even after fixation.

### Photostability of fluorescent protein tags

The photostability of fluorescent protein tags plays a critical role in time-lapse experiments, such as the measurement of intraflagellar transport rates or the trafficking of secreted cargo. To determine the half-lives of fluorescent proteins under experimentally relevant conditions in live cells, a 30 × 1-second time-lapse movie was conducted on each cell line where a spot corresponding to the tagged TZP157 was visible.

While mCherry and mCardinal were found to be the most photostable, their dimness limits their utility in most experimental setups (Figure 3). In contrast, 3xmNG and mNG were found to exhibit the best combination of brightness and stability, remaining the brightest protein tags throughout the 30-second time-lapse. In this example, the size of the triple tandem repeat did not interfere with protein localization, and in our experience, proteins such as IFT components tolerate a tag of this size well. However, in experiments where the size of the protein tag is a concern, a single mNeonGreen tag may be more suitable. Amongst the red fluorescent proteins, although tdTomato was found to be the brightest, tagRFPt [16] exhibited the optimal combination of brightness and photostability and was the brightest tag after 24 seconds of exposure.

### mScarlet slow maturation causes an experimental artifact in tagging experiments

The pPOT vector series, with its inherent diversity, enables the generation of intricate cell lines for co-tagging experiments. Using this technique, we tagged two proteins, TZP103.8 and basalin, that play crucial roles in nucleating the flagellum central pair of microtubules that coordinate flagellar dynein activity [13,17].

Our observations indicated that TZP103.8 tagged with mScarlet was evident only in the late stages of flagellum maturation, after the central pair had been assembled and inconsistent with its known role in central pair assembly (Figure 4a). Indeed, when examining cytoskeletons in which a new flagellum was visible by phase contrast microscopy but the mitochondrial DNA (kinetoplast) had not yet segregated, we noted that more than 75% of the assembling flagella were negative for mScarlet-tagged TZP103.8 (Figure 4b).

**Figure 4:**
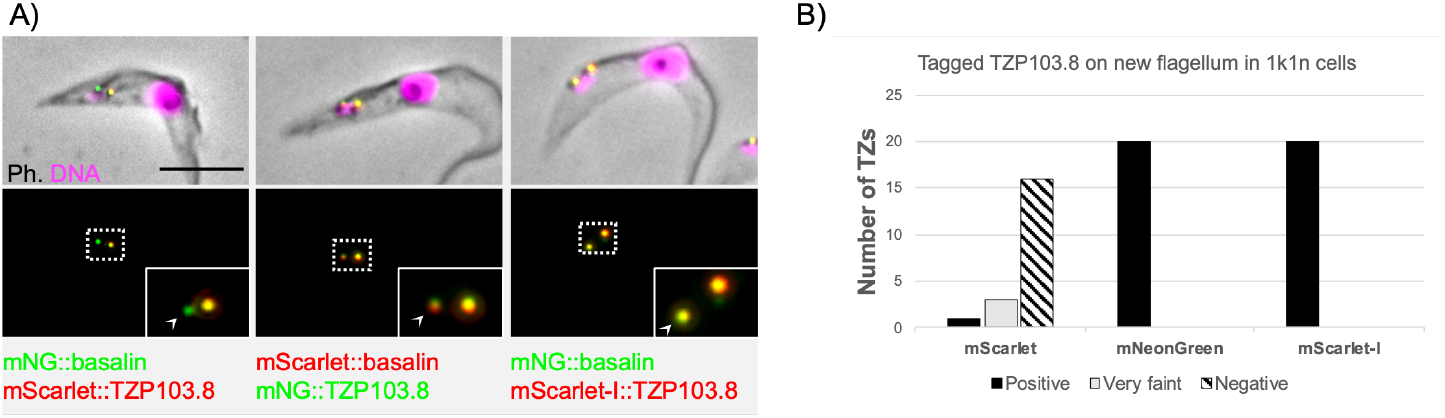
Absence of mScarlet::TZP103.8 from assembling flagella is an experimental artifact. A) Illustrative micrographs of basalin and TZP103.8 tagged with different fluorescent proteins to examine their assembly. When TZP103.8 was tagged with mScarlet (left) it was not evident on >75% of flagella during the earliest stages of biogenesis. In contrast, TZP103.8 is detected on all new flagella when tagged with mNeonGreen (middle) or the fast-folding variant, mScarlet-I (right). Arrowheads indicate newly assembling flagella in dividing cells, Scale bar = 5 *μ*m. B) Twenty dividing cells in the very early stages of flagellar assembly were assessed for the presence of tagged TZP103.8 on the new flagellum. 1k1n refers to the number of kinetoplast (k) and nuclei (n) and indicates very early stages of flagellum assembly.

To investigate this further, we swapped the tags, tagging TZP103.8 with mNeonGreen and basalin with mScarlet. With this change, all new flagella were positive for tagged TZP103.8. A review of the literature suggested that the slow maturation of mScarlet may have caused this apparent absence [6,12,18]. To test this hypothesis, we employed mScarlet-I, a fast-folding variant of mScarlet [6,19] that differs in a single amino acid change proximal to the chromophore. When we used mScarlet-I as a tag, TZP103.8 was detectable at the very earliest stages of flagellum assembly, consistent with mNeonGreen (Figure 4).

Our findings indicate that mScarlet-I is a more suitable protein tag in most experimental setups, and that caution should be exercised when using fluorescent proteins as protein tags. Nevertheless, the delayed onset of mScarlet fluorescence may serve as a “molecular timer” to determine the relative age of different cellular pools of protein.

## Discussion

The pPOT plasmids described here signify a notable improvement on the previous generation of pPOT vectors [3]. These new plasmids have a range of tags and drug resistances that allow for the creation of cell lines with multiple different tagged proteins. This enables co-localisation and biochemistry studies to be conducted. Moreover, the use of universal primer annealing sequences considerably reduces the cost and enhances the flexibility of the system because a single long-primer pair can be used to tag a target protein using a variety of protein tags or drug resistances (Supplemental File 3).

The modularity of these vectors allows for the easy incorporation of new primary tags and flanking epitopes into the toolkit as improved biochemical or fluorescent tags become available. Additionally, the development of modular “fusion tags” makes it possible to tag proteins with tags that have multiple functions. This enables the combination of, for instance, bioluminescence and fluorescence imaging, as elegantly demonstrated in *Trypanosoma cruzi* [20], to locate parasites more easily in infected tissue. Similarly, other fusion tags described here, which combine fluorescent tags with proximity-dependent biotinylation, enable the confirmation of localisation and the subsequent identification of interacting proteins. The development of a set of vectors that incorporate ten copies of five different epitope tags supports the emerging technique of expansion microscopy because it facilitates tagging of multiple proteins for ultrastructural protein cartography. Moreover, split fluorescent protein tags, click chemistry (HaloTag, CLIP and SNAP tags) and proximity-dependant biotinylation (BirA* and BIOID2) tags support easy and scalable protein-protein interaction studies.

The pPOT plasmids discussed here have not been tested in related trypanosomatids, such as *T. congolense* or *Leishmania mexicana*. However, kinetoplastids appear to accept foreign regulatory nucleotide sequences [21], suggesting they could potentially function in these cells, although CRISPR technology might be required to enhance integration efficiency (Beneke et al., 2017; Kovářová et al., 2022). In fact, the incorporation of exogenous DNA might even offer an advantage by reducing interference with the organisms’ endogenous gene regulation.

We tested the utility of 17 fluorescent proteins in microscopy imaging and showed that mNeonGreen remains the brightest fluorescent protein even after extended illumination and fixation, making it the best choice in most applications. This is supported by its success in the TrypTag whole-genome tagging project [22,23] and in tagging a variety of cytological markers [24]. We also observed a putative “slow folding” artifact of mScarlet, meaning that, for most purposes, the fast-folding variant mScarlet-I is the preferred red fluorescent protein. However, for time-lapse imaging, tagRFPt would be the preferred red fluorescent protein, as it performs well over a 30-second time-lapse movie.

High-intensity irradiation using a metal-halide lamp and standard filter sets were used in this study. Optimisation of such parameters would likely improve performance. Nonetheless, our data underscores that the behaviour of fluorescent proteins differs as protein tags inside a living cell from that predicted by their in vitro determined characteristics, such as quantum yield. Moreover, it is likely that fluorescent proteins perform differently in different cell types or different cellular locations [12]. Therefore, testing protein tags in a realistic setting is essential.

We anticipate that this extensive set of more than one hundred pPOT vectors will be an important toolset for investigating trypanosome protein localisation and function.

## Materials and methods

### Microscopy

Images were acquired using a Leica DM6 B microscope equipped with a metal halide lamp (EL 6000, 11504115) serving as the illumination source, a 63x objective with a numerical aperture of 1.30 (11506385) and a Leica K5 Microscope Camera (11547112). Filters used for fluorescence imaging depended on the spectral properties of the fluorescent proteins: blue fluorescent proteins: DAPI ET, 11504203; green fluorescent proteins: GFP ET, 11504164; red fluorescent proteins: RHOD ET, 11504205; and far-red fluorescent protein: Y5 ET, 11504171.

Samples were prepared for live-cell and fixed-cell imaging as described [25]. Briefly, cells were washed and settled on clean glass slides prior to imaging (live-cell) or fixation. Cells were either fixed in cold methanol or 2% formaldehyde for 10 minutes and fixed cells were mounted in phosphate-buffered glycerol containing DABCO and imaged immediately.

Data for calculating brightness and photostability of live cells was acquired by performing a 30 × 1 second time lapse, with the first image in the series being used to calculate brightness, and the entire series being used to calculate photostability. Data to calculate the brightness of fluorescent proteins under different fixation conditions was acquired by performing a single 2-second exposure.

### Image analysis

Spots corresponding to tagged TZP157 were initially identified using FIJI’s built-in “Find Maxima” and then fields were manually curated. Brightness of each spot was quantified using a custom FIJI script (Supplemental files 1 and 2). Briefly, a square was drawn around each maxima and the signal intensity measured, and the local background subtracted using the median pixel value of a larger square centred on the TZP157 spot. This was repeated on each spot corresponding to tagged TZP157 at each timepoint, with more than 100 cells analysed per timepoint. The photostability half-life of each protein was determined by calculating the linear interpolation between the two time points that were closest to half the brightness value after one second.

### Plasmids and cell-lines

Modules were cloned into the pPOT vector using standard molecular biology techniques. Where necessary, genes were synthesized using either Twist Bioscience or Life technologies, with custom scripts performing codon optimisation based on the codon frequency of the trypanosome genome. Primers for gene tagging were designed using TagIt [3] and primer designs for tagging every Trypanosome gene in the 927 and 427 reference genomes are provided in Supplemental File 3 as a community resource. Amplicons and transgenic cell lines were produced as described [3]. Procyclic form TREU927 reference genome cells were used for all experiments.

## Acknowledgements and contributions

AP performed experiments and edited the manuscript. SD designed the project, wrote the manuscript, cloned plasmids, performed experiments and analysed the data. JT and JDS cloned plasmids. SV, KG and SD provided funding and supervision. We would like to thank Martin Tylor for providing the Ppyr9eh gene, Giridhar Chandrasekharan for the BIOID2 gene, and Division of Biomedical Sciences Warwick Medical School for their generous support. The authors declare that they have no competing interests.

## Supplemental Figures

**Supplementary Figure 1.**
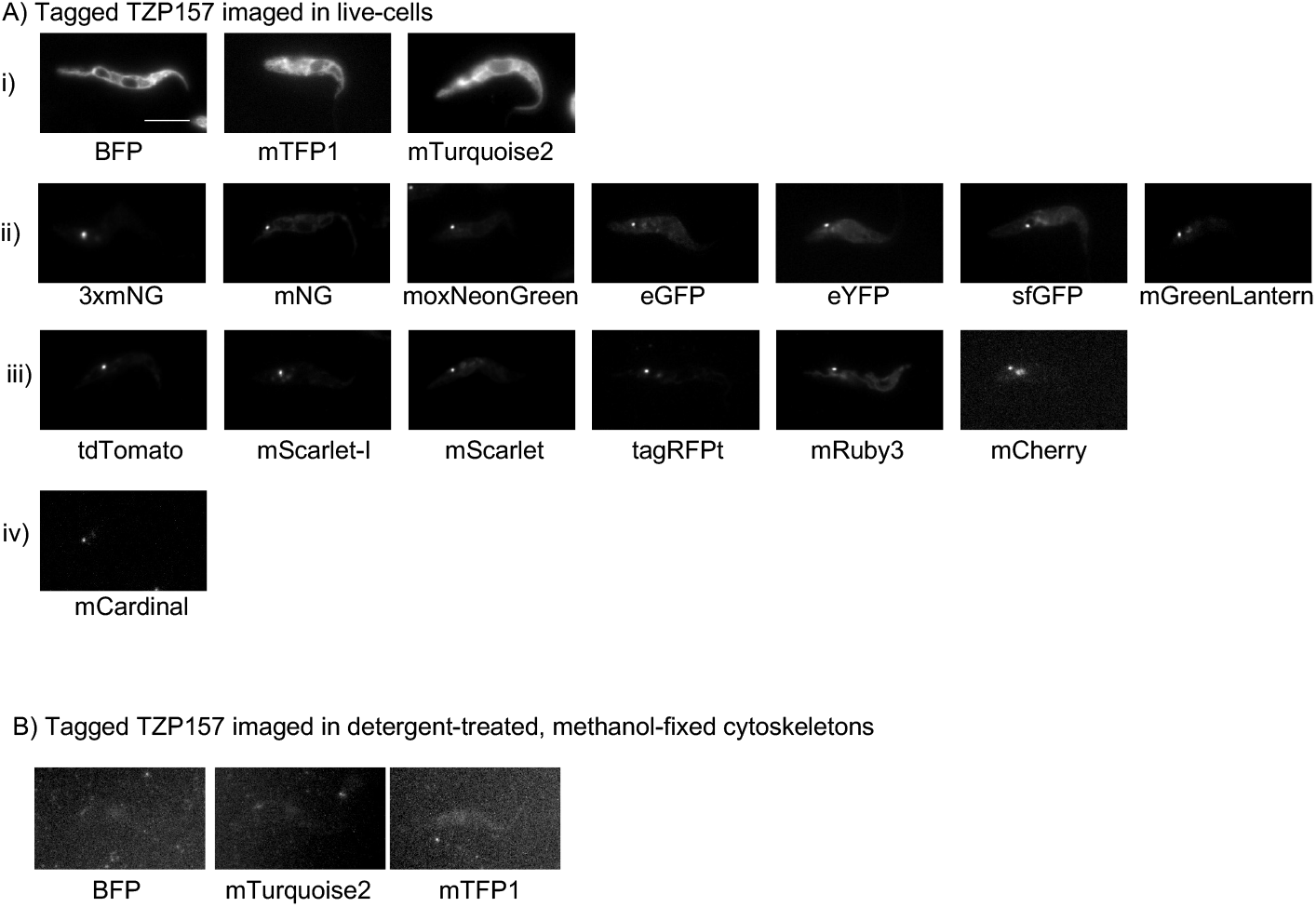
Examples micrograph for each fluorescent protein tested as a protein tag in this study. A) Live cells expressing tagged TZP157 tagged with (i) blue, (ii) green, (iii) red and (iv) far-red fluorescent proteins. B) Cells expressing TZP157 tagged with BFP, mTurquoise2 and mTFP1 were treated with detergent to reduce auto-fluorescent background from the cytosol. Scale bar = 5 *μ*m.

**Supplementary Figure 2.**
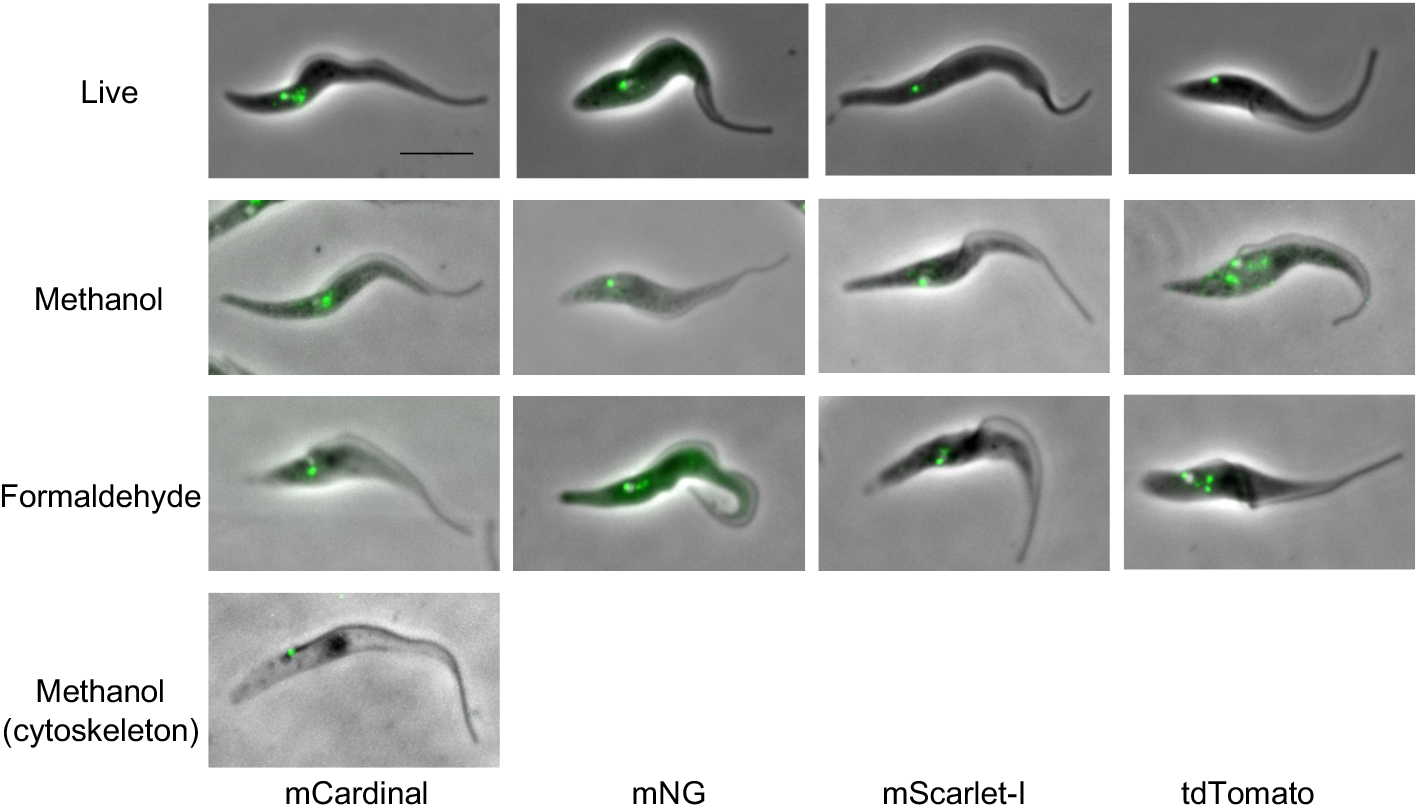
Effect of fixation upon the brightness of fluorescent proteins. Cells expressing TZP157 tagged with mCardinal, mNeonGreen, mScarlet-I and tdTomato were imaged by fluorescence microscopy after fixation with either methanol or formaldehyde. mCardinal::TZP157 was not visible in fixed whole cells but was detected after extraction of soluble material. Scale bar = 5 *μ*m.

**Supplemental Table 1**

A list of the pPOT vectors currently on Addgene for distribution to the community.

**Supplemental File 1**

A custom FIJI script to quantify the brightness and photostability of tagged TZP157 over a 30-second time-course

**Supplemental File 2**

A custom FIJI script to quantify the brightness of tagged TZP157 in live cells and fixed cells.

**Supplemental File 3**

Primer sequences for tagging every protein encoded in the 427 and 927 trypanosome reference genomes on the N and C terminus.

