## Supplemental Table 1 for "Unlocking Trypanosome Biology: A Comprehensive Protein-Tagging Toolkit for Localization and Functional Analysis"

| Addgene ID | Name | Drug resistance | Primary tag | Secondary Tag |
|---|---|---|---|---|
| 181940 | pPOTv7-blast-10FLAG | blast | 10x FLAG | none |
| 181941 | pPOTv7-blast-10HA | blast | 10x HA | None |
| 181939 | pPOTv7-blast-10Myc | blast | 10x Myc | None |
| 179807 | pPOTv7-blast-10Ty | blast | 10x Ty | None |
| 181938 | pPOTv7-blast-10V5 | blast | 10x V5 | None |
| 179827 | pPOTv7-blast-3HaloTag | blast | 3x HaloTag | None |
| 179831 | pPOTv7-blast-3mNG | blast | 3x mNG | None |
| 190258 | pPOTv7-blast-BFP | blast | BFP | None |
| 179788 | pPOTv7-blast-birA* | blast | birA | None |
| 179787 | pPOTv7-blast-CLIPtag | blast | CLIP | None |
| 179781 | pPOTv7-blast-eGFP | blast | eGFP | None |
| 179784 | pPOTv7-blast-eYFP | blast | eYFP | None |
| 179795 | pPOTv7-blast-lssmOrange | blast | lss mOrange | None |
| 179796 | pPOTv7-blast-mCardinal | blast | mCardinal | None |
| 179789 | pPOTv7-blast-mCherry | blast | mCherry | None |
| 190221 | pPOTv7-blast-mGL | blast | mGreenLantern | None |
| 179817 | pPOTv7-blast-mNG | blast | mNG | None |
| 179801 | pPOTv6-blast-3Myc:mNG:3Myc | blast | mNG | c-Myc |
| 201094 | pPOTv6-blast-His:mNG:His | blast | mNG | His |
| 179783 | pPOTv6-blast-3Ty:mNG:3Ty | blast | mNG | Ty1 |
| 179785 | pPOTv7-blast-mNG:HaloTag | blast | mNG::tbHaloTag | None |
| 201441 | pPOTv7-blast-mNG:Ppyr9eh | blast | mNG:Ppyr9eh | None |
| 190209 | pPOTv7-blast-mNG2(1-10) | blast | mNG2(1-10) | None |
| 190210 | pPOTv7-blast-mNG2(11) | blast | mNG2(11) | None |
| 190222 | pPOTv7-blast-moxNG | blast | moxNG | None |
| 179793 | pPOTv7-blast-mRuby3 | blast | mRuby3 | None |
| 179818 | pPOTv7-blast-mScarlet | blast | mScarlet | None |
| 201442 | pPOTv7-blast-mScarletI | blast | mScarlet-I | None |
| 179802 | pPOTv6-blast-3Myc:mScarletI:3Myc | blast | mScarlet-I | c-Myc |
| 201097 | pPOTv6-blast-His:mScarletI:His | blast | mScarlet-I | His |
| 201090 | pPOTv6-blast-3Ty:mScarletI:3Ty | blast | mScarlet-I | Ty1 |
| 179792 | pPOTv7-blast-mTFP1 | blast | mTFP1 | None |
| 179791 | pPOTv7-blast-mTurquoise2 | blast | mTurquoise2 | None |
| 179797 | pPOTv7-blast-nanoluc | blast | nanoluc | None |
| 179824 | pPOTv6-blast-3Ty:pHlourinM153R:3Ty | blast | pHlourin M153R | Ty1 |
| 179806 | pPOTv6-blast-3Ty:pHlourin2:3Ty | blast | pHlourin2 | Ty1 |
| 201443 | pPOTv7-blast-Ppyr9eh | blast | Ppyr9eh | None |
| 190206 | pPOTv7-blast-sfCherry2(1-10) | blast | sfCherry2(1-10) | None |
| 190207 | pPOTv7-blast-sfCherry2(11) | blast | sfCherry2(11) | None |
| 179782 | pPOTv7-blast-sfGFP | blast | sfGFP | None |
| 179786 | pPOTv7-blast-SNAPtag | blast | SNAP | None |
| 190208 | pPOTv7-blast-spGFP(1-10) | blast | spGFP(1-10) | None |
| 179805 | pPOTv6-blast-3Ty:syPher2:3Ty | blast | syPher2 | Ty1 |
| 179790 | pPOTv7-blast-tagRFPt | blast | tagRFPt | None |
| 201086 | pPOTv6-blast-3Ty:tagRFPt:3Ty | blast | tagRFPt | Ty1 |
| 181953 | pPOTv7-blast-HaloTag | blast | tbHaloTag | None |
| 181943 | pPOTv6-blast-3Myc:HaloTag:3Myc | blast | tbHaloTag | c-Myc |
| 181942 | pPOTv6-blast-3Ty:HaloTag:3Ty | blast | tbHaloTag | Ty1 |
| 179794 | pPOTv7-blast-tdTomato | blast | tdTomato | None |
| 179812 | pPOTv7-g418-10FLAG | g418 | 10x FLAG | None |
| 179813 | pPOTv7-g418-10HA | g418 | 10x HA | None |
| 179811 | pPOTv7-g418-10Myc | g418 | 10x Myc | None |
| 179810 | pPOTv7-g418-10V5 | g418 | 10x V5 | None |
| 179829 | pPOTv7-g418-3HaloTag | g418 | 3x HaloTag | None |
| 179832 | pPOTv7-g418-3mNG | g418 | 3x mNG | None |
| 190259 | pPOTv7-g418-BFP | g418 | BFP | None |
| 190220 | pPOTv7-g418-eYFP | g418 | eYFP | None |
| 190261 | pPOTv7-g418-mCherry | g418 | mCherry | None |
| 179821 | pPOTv7-g418-mNG | g418 | mNG | None |
| 190217 | pPOTv6-g418-3Myc-mNG-3Myc | g418 | mNG | c-Myc |
| 201095 | pPOTv6-g418-His:mNG:His | g418 | mNG | His |
| 179800 | pPOTv6-g418-3Ty:mNG:3Ty | g418 | mNG | Ty1 |
| 190214 | pPOTv7-g418-mNG2(1-10) | g418 | mNG2(1-10) | None |
| 190215 | pPOTv7-g418-mNG2(11) | g418 | mNG2(11) | None |
| 179823 | pPOTv7-g418-mScarlet | g418 | mScarlet | None |
| 179822 | pPOTv7-g418-mScarletI | g418 | mScarlet-I | None |
| 201098 | pPOTv6-g418-His:mScarletI:His | g418 | mScarlet-I | His |
| 201091 | pPOTv6-g418-3Ty:mScarletI:3Ty | g418 | mScarlet-I | Ty1 |
| 190211 | pPOTv7-g418-sfCherry2(1-10) | g418 | sfCherry2(1-10) | None |
| 190212 | pPOTv7-g418-sfCherry2(11) | g418 | sfCherry2(11) | None |
| 190213 | pPOTv7-g418-spGFP(1-10) | g418 | spGFP(1-10) | None |
| 190257 | pPOTv7-g418-spGFP(11) | g418 | spGFP11 | None |
| 201085 | pPOTv7-g418-tagRFPt | g418 | tagRFPt | None |
| 201087 | pPOTv6-g418-3Ty:tagRFPt:3Ty | g418 | tagRFPt | Ty1 |
| 179815 | pPOTv7-hygro-10FLAG | hygro | 10x FLAG | None |
| 179816 | pPOTv7-hygro-10HA | hygro | 10x HA | None |
| 179814 | pPOTv7-hygro-10Myc | hygro | 10x Myc | None |
| 179808 | pPOTv7-hygro-10Ty | hygro | 10x Ty | None |
| 179809 | pPOTv7-hygro-10V5 | hygro | 10x V5 | None |
| 179828 | pPOTv7-hygro-3HaloTag | hygro | 3x HaloTag | None |
| 179833 | pPOTv7-hygro-3mNG | hygro | 3x mNG | None |
| 190260 | pPOTv7-hygro-BFP | hygro | BFP | None |
| 190219 | pPOTv7-hygro-eYFP | hygro | eYFP | None |
| 190262 | pPOTv7-hygro-mCherry | hygro | mCherry | None |
| 179819 | pPOTv7-hygro-mNG | hygro | mNG | None |
| 190218 | pPOTv6-hygro-3Myc-mNG-3Myc | hygro | mNG | c-Myc |
| 201096 | pPOTv6-hygro-His:mNG:His | hygro | mNG | His |
| 179799 | pPOTv6-hygro-3Ty:mNG:3Ty | hygro | mNG | Ty1 |
| 190263 | pPOTv7-hygro-mScarlet | hygro | mScarlet | None |
| 179820 | pPOTv7-hygro-mScarletI | hygro | mScarlet-I | None |
| 201099 | pPOTv6-hygro-His:mScarletI:His | hygro | mScarlet-I | His |
| 201092 | pPOTv6-hygro-3Ty:mScarletI:3Ty | hygro | mScarlet-I | Ty1 |
| 190216 | pPOTv7-hygro-mNG2(1-10) | hygro | spNG2(1-10) | None |
| 201084 | pPOTv7-hygro-tagRFPt | hygro | tagRFPt | None |
| 201088 | pPOTv6-hygro-3Ty:tagRFPt:3Ty | hygro | tagRFPt | Ty1 |
| 181935 | pPOTv7.1-phleo-mNG | phleo | mNG | None |
| 181937 | pPOTv6.1-phleo-3Myc:mNG:3Myc | phleo | mNG | c-Myc |
| 181936 | pPOTv6.1-phleo-Ty:mNG:Ty | phleo | mNG | Ty1 |
| 179803 | pPOTv6-puro-3Myc:mNG:3Myc | puro | mNG | c-Myc |
| 179798 | pPOTv6-puro-3Ty:mNG:3Ty | puro | mNG | Ty1 |
| 179804 | pPOTv6-puro-3Myc:mScarletI:3Myc | puro | mScarlet-I | c-Myc |
| 201093 | pPOTv6-puro-3Ty:mScarletI:3Ty | puro | mScarlet-I | Ty1 |
| 201089 | pPOTv6-puro-3Ty:tagRFPt:3Ty | puro | tagRFPt | Ty1 |
